# Contracted Functional Connectivity Profiles in Autism

**DOI:** 10.1101/2024.02.21.581260

**Authors:** Clara F. Weber, Valeria Kebets, Oualid Benkarim, Sara Lariviere, Yezhou Wang, Alexander Ngo, Hongxiu Jiang, Xiaoqian Chai, Bo-yong Park, Michael P. Milham, Adriana Di Martino, Sofie Valk, Seok-Jun Hong, Boris C. Bernhardt

## Abstract

**Objective:** Autism spectrum disorder (ASD) is a pervasive neurodevelopmental condition that is associated with atypical brain network organization, with prior work suggesting differential connectivity alterations with respect to functional connection length. Here, we tested whether functional connectopathy in ASD specifically relates to disruptions in long-relative to short-range functional connectivity profiles. Our approach combined functional connectomics with geodesic distance mapping, and we studied associations to macroscale networks, microarchitectural patterns, as well as socio-demographic and clinical phenotypes.

**Methods:** We studied 211 males from three sites of the ABIDE-I dataset comprising 103 participants with an ASD diagnosis (mean±SD age=20.8±8.1 years) and 108 neurotypical controls (NT, 19.2±7.2 years). For each participant, we computed cortex-wide connectivity distance (CD) measures by combining geodesic distance mapping with resting-state functional connectivity profiling. We compared CD between ASD and NT participants using surface-based linear models, and studied associations with age, symptom severity, and intelligence scores. We contextualized CD alterations relative to canonical networks and explored spatial associations with functional and microstructural cortical gradients as well as cytoarchitectonic cortical types.

**Results:** Compared to NT, ASD participants presented with widespread reductions in CD, generally indicating shorter average connection length and thus suggesting reduced long-range connectivity but increased short-range connections. Peak reductions were localized in transmodal systems (*i*.*e*., heteromodal and paralimbic regions in the prefrontal, temporal, and parietal and temporo-parieto-occipital cortex), and effect sizes correlated with the sensory-transmodal gradient of brain function. ASD-related CD reductions appeared consistent across inter-individual differences in age and symptom severity, and we observed a positive correlation of CD to IQ scores.

**Conclusions:** Our study showed reductions in CD as a relatively stable imaging phenotype of ASD that preferentially impacted paralimbic and heteromodal association systems. CD reductions in ASD corroborate previous reports of ASD-related imbalance between short-range overconnectivity and long-range underconnectivity.

## INTRODUCTION

Autism spectrum disorder (ASD) is a prevalent and pervasive neurodevelopmental condition [1, 2], commonly manifesting in atypical social cognition and communication, repetitive behaviors and interests, sometimes together with imbalances in affective, sensory, and perceptual processing [2-4]. Despite extensive research, pathomechanisms of ASD remain incompletely understood. Convergent evidence from molecular, histological, and neuroimaging work suggests atypical brain network organization, motivating continued efforts to identify substrates of autism connectopathy [5-10].

By interrogating brain structure and function *in vivo*, magnetic resonance imaging (MRI) lends itself as a window into the human connectome [11, 12]. Resting-state functional MRI (rs-fMRI) [13-16] can probe whole-brain intrinsic functional networks [17-20], both in terms of functionality and spatial layout [21-25]. Moreover, rs-fMRI analysis has become common in the study of typical and atypical neurodevelopment [5, 25, 26], and in identifying substrates of symptom profiles in complex neurodevelopmental conditions such as ASD [25, 27-30]. Across the cortex, there is an important inter-regional variability of functional connection length: patterns of strong local connectivity are typical for unimodal areas, including the somatosensory and visual systems, where fast and efficient local signal transmission is necessary [23, 31]. Contrarily, long-range connections are increasingly found in heteromodal association systems and paralimbic networks, systems that implicate more integrative, cognitive-related processing [31, 32]. While longer connections are metabolically expensive to build and maintain [31, 33], they conversely provide gains in terms of processing flexibility and integrative capacity [23, 31, 32].

Increasing evidence suggests widespread functional connectivity alterations in ASD, generally reporting mosaic patterns of under- and overconnectivity in ASD as compared to neurotypical controls (NT) [5, 34-37]. In individuals with ASD, short- and long-range connectivity is likely affected differently [34, 38]. While underconnectivity is reported at a global level and in transmodal systems, such as the default mode network (DMN) in ASD [5, 35, 39], there is some literature also emphasizing overconnectivity, primarily in unimodal cortical areas and subcortico-cortical networks [9, 35, 40]. It has been hypothesized that ASD is associated with long-range underconnectivity, but short-range overconnectivity [38, 41, 42]. However, as studies report diverging results, existing evidence fails to provide a comprehensive and mechanistic explanation and spatial mapping of ASD-related connectivity alterations. Functional connectivity and distance length are spatially linked, as long- and short-range connections are neither randomly nor evenly distributed across the cortex, but characteristic connection length of a cortical region mirrors its position in the putative cortical hierarchy [31, 32]. Thus, combining functional neuroimaging and topological information can provide further insights into connectivity shifts in ASD. Previous studies have proposed connectivity distance (CD) as a metric that combines functional connectivity with measures of spatial proximity between brain areas [31, 38, 43], notably by calculating geodesic distance along the cortical surface [43, 44]. CD is the average distance to connected nodes of a given vertex, thus capturing functionally relevant connection lengths [31, 43]. In NT populations, CD has been found to increase with distance to primary cortical areas [31], supporting that short-range connections predominate in unimodal regions while long-range connections are increasingly present in transmodal association systems, such as the DMN. Additional evidence has underlined impaired segregation and integration between cortical hierarchies in ASD mirrored in functional neuroimaging [45].

The well-recognized etiological heterogeneity in ASD motivates contextualization of neuroimaging-based phenotypes against established measures of neural organization, to explore potential pathways of susceptibility. Converging evidence hints towards a relationship between CD and cortical microarchitecture [31, 32, 43, 46, 47], notably cellular composition, columnar topography, and lamination of the cortex [46, 48]. Local increases in cell density and smaller pyramidal cells [49-51], suggestive of short-range overconnectivity at the expense of long-range connections, could potentially impact macrolevel connections and circuit function [50, 52]. Neuronal cell size, density, and connection types vary across cortical areas and their modalities, as observed in histological studies as early as in the foundational descriptions of cortical types by Von Economo and Koskinas [53]. The microstructural organization of the cortex is generally thought to correlate with large-scale functional organization, with unimodal regions exhibiting stronger lamination patterns while transmodal systems express reduced laminar differentiation [54]. More recently, analyzing *post mortem* histological reconstructions of the human brain has allowed for additional histological contextualization of imaging findings [55-59]. In particular, the 3D BigBrain dataset has been used to derive gradients of microstructural cortical organization that run from sensory to paralimbic systems [60]. Despite some notable differences [60, 61], this hierarchical axis is in most parts converging with foundational descriptions of the functional cortical hierarchy [23], and decompositions of resting-state functional connectivity [24, 62, 63]. As such, these novel resources set the stage to explore whether connectome contractions in ASD follow microstructural and functional gradients, indicating a connectopathological susceptibility that follows established principles of cortical hierarchical organization.

Considering the mounting evidence of the differential impact of long-*vs* short-range connectivity alterations in ASD, we aimed to investigate CD in individuals with ASD and NT as a possible underpinning of shorter average connection length, *i*.*e*. contracted connectome profiles in ASD. We leveraged the multi-centric Autism Brain Imaging Data Exchange (ABIDE-I) repository [64] and quantified CD alterations in ASD relative to NT by combining rs-fMRI connectivity analysis with cortex-wide geodesic distance mapping. We contextualized ASD-related CD reductions across functional networks and sensory-transmodal gradients of cortical functional hierarchy [24]. On a microscopic scale, we investigated correlation to histology-derived gradient maps and cortical types, to find a more complete explanatory model for connectome contractions in ASD.

## METHODS

### Participants

We studied data from the first release of the Autism Brain Imaging Data Exchange (ABIDE) [64], a multi-centric data and imaging collection consortium. Similar to previous work [25, 65, 66], we included imaging and clinical data from three different sites that included data from >=10 ASD and NT respectively, *i*.*e*., University of Pittsburgh School of Medicine (PITT), New York University Langone Medical Center (NYU), and University of Utah School of Medicine (USM). ABIDE data has been collected in alignment with local institutional review board frameworks and made publicly available in an anonymized form in accordance with the Health Insurance Portability and Accountability Act (HIPAA) guidelines. ASD individuals were identified in an inperson diagnostic interview using the Autism Diagnostic Observation Schedule (ADOS), and subjects with genetic disorders associated to ASD, or contraindications to scanning such as pregnancy were excluded from the study. NT controls did not have any history of psychiatric disease. Due to the small number of female participants included in the first ABIDE release [64], we limited our analysis to male individuals. Additionally, we only retained cases with acceptable T1 imaging quality and surface extraction outcomes. After excluding cases with high head motion as described below, we obtained a sample size of n=211 (103/108 ASD/NT). To assess symptom severity, we considered ADOS scores [4], which evaluate three characteristic symptomatic domains of ASD, *i*.*e*., communication and language, reciprocal social interactions and restricted/repetitive behaviors. Furthermore, we considered intelligence quotient (IQ) and IQ subscores for verbal and performance IQ as measured by the Wechsler Adult Intelligence Scale [67]. Since previous literature suggests cognitive imbalances, such as high variability between verbal and nonverbal abilities, in ASD [68, 69], we additionally calculated and analyzed the ratio of verbal over nonverbal IQ [70]. Detailed demographic information is provided in **Table 1**.

**Table 1.**
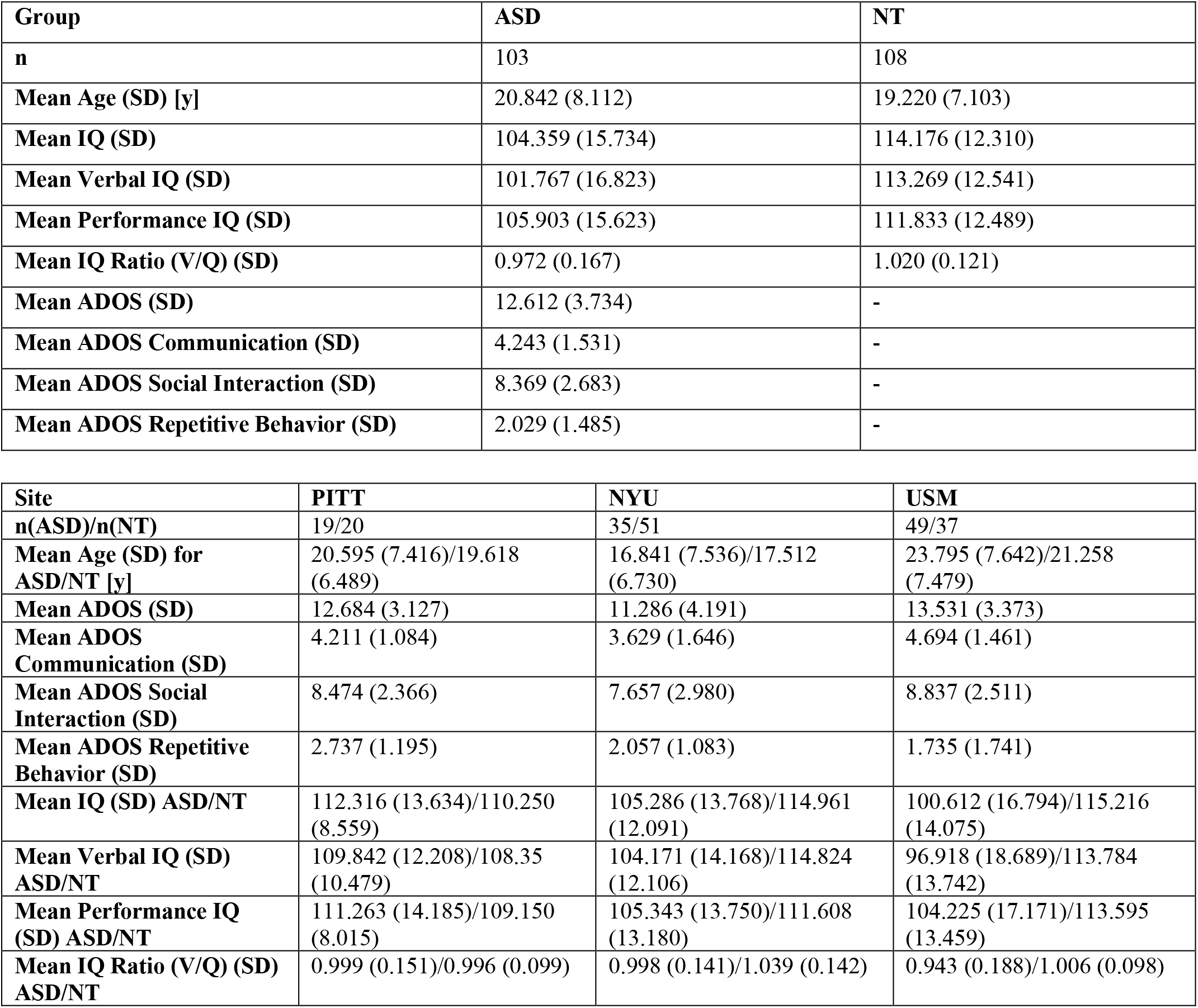
Study cohort demographic information for ASD and neurotypical control (NT) groups. Age in years (y), SD = standard deviation.

| Group | ASD | NT |
| --- | --- | --- |
| **n** | 103 | 108 |
| **Mean Age (SD) [y]** | 20.842 (8.112) | 19.220 (7.103) |
| **Mean IQ (SD)** | 104.359 (15.734) | 114.176 (12.310) |
| **Mean Verbal IQ (SD)** | 101.767 (16.823) | 113.269 (12.541) |
| **Mean Performance IQ (SD)** | 105.903 (15.623) | 111.833 (12.489) |
| **Mean IQ Ratio (V/Q) (SD)** | 0.972 (0.167) | 1.020 (0.121) |
| **Mean ADOS (SD)** | 12.612 (3.734) | - |
| **Mean ADOS Communication (SD)** | 4.243 (1.531) | - |
| **Mean ADOS Social Interaction (SD)** | 8.369 (2.683) | - |
| **Mean ADOS Repetitive Behavior (SD)** | 2.029 (1.485) | - |

| Site | PITT | NYU | USM |
| --- | --- | --- | --- |
| **n(ASD)/n(NT)** | 19/20 | 35/51 | 49/37 |
| **Mean Age (SD) for ASD/NT [y]** | 20.595 (7.416)/19.618 (6.489) | 16.841 (7.536)/17.512 (6.730) | 23.795 (7.642)/21.258 (7.479) |
| **Mean ADOS (SD)** | 12.684 (3.127) | 11.286 (4.191) | 13.531 (3.373) |
| **Mean ADOS Communication (SD)** | 4.211 (1.084) | 3.629 (1.646) | 4.694 (1.461) |
| **Mean ADOS Social Interaction (SD)** | 8.474 (2.366) | 7.657 (2.980) | 8.837 (2.511) |
| **Mean ADOS Repetitive Behavior (SD)** | 2.737 (1.195) | 2.057 (1.083) | 1.735 (1.741) |
| **Mean IQ (SD) ASD/NT** | 112.316 (13.634)/110.250 (8.559) | 105.286 (13.768)/114.961 (12.091) | 100.612 (16.794)/115.216 (14.075) |
| **Mean Verbal IQ (SD) ASD/NT** | 109.842 (12.208)/108.35 (10.479) | 104.171 (14.168)/114.824 (12.106) | 96.918 (18.689)/113.784 (13.742) |
| **Mean Performance IQ (SD) ASD/NT** | 111.263 (14.185)/109.150 (8.015) | 105.343 (13.750)/111.608 (13.180) | 104.225 (17.171)/113.595 (13.459) |
| **Mean IQ Ratio (V/Q) (SD) ASD/NT** | 0.999 (0.151)/0.996 (0.099) | 0.998 (0.141)/1.039 (0.142) | 0.943 (0.188)/1.006 (0.098) |

### MRI Acquisition Parameters

Data from all three sites were acquired on 3T Siemens scanners. Acquisition protocols for T1-weighted (T1w) and rs-fMRI were as follows for the three included sites: *(i) NYU*. Data were acquired on Allegra scanner using 3D-TurboFLASH for T1w (repetition time (TR) = 2530 ms; echo time (TE) = 3.25 ms; inversion time (TI) = 1100 ms; flip angle = 7**°**; matrix = 256 × 256; 1.3 × 1.0 × 1.3 mm^3^ voxels) and 2D-echo planar imaging (EPI) for rs-fMRI (TR = 2000ms; TE = 15 ms; flip angle = 90**°**; matrix = 80 × 80; 180 volumes, 3.0 × 3.0 × 4.0 mm^3^ voxels); *(ii) PITT*. Data were acquired on an Allegra scanner using 3D-MPRAGE for T1w (TR = 2100 ms; TE = 3.93 ms; TI = 1000 ms; flip angle = 7**°**; matrix = 269 × 269; 1.1 × 1.1 × 1.1 mm^3^ voxels) and 2D-EPI for rs-fMRI (TR = 1500 ms; TE = 35 ms; flip angle = 70**°**; matrix = 64 × 64; 200 volumes, 3.1 × 3.1 × 4.0 mm^3^ voxels); *(iii) USM*. Data were acquired on a TrioTim scanner using 3D-MPRAGE for T1w (TR = 2300 ms; TE = 2.91 ms; TI = 900 ms; flip angle = 9**°**; matrix = 240 × 256; 1.0 × 1.0 × 1.2 mm^3^ voxels) and 2D-EPI for rs-fMRI (TR = 2000ms; TE = 28 ms; flip angle = 90**°**; matrix = 64 × 64; 240 volumes; 3.4 × 3.4 × 3.0 mm^3^ voxels).

### MRI Processing

#### a) Structural MRI processing

1. T1w data were preprocessed using FreeSurfer v5.1 [71-73] (https://surfer.nmr.mgh.harvard.edu), which included bias field correction, intensity normalization, removal of non-brain tissue, and white matter segmentation. For a more accurate gray/white matter separation and extraction of a vertex-based surface, a mesh model was fit onto the white matter volume. Resulting surfaces were subsequently brought into spherical representation to improve alignment to sulco-gyral patterns.

#### b) Geodesic distance

We calculated geodesic distance as the physical distance of vertices along the pial surface. Specifically, we computed each individual’s intra-hemispheric geodesic distance map between all pairs of vertices within each hemisphere, using the HCP Workbench *surface-geodesic-distance* command [74] (https://www.humanconnectome.org/software/workbench-command) and resampled data to Conte69 surface space to obtain 10,242 vertices per hemisphere [75] (https://github.com/Washington-University/HCPpipelines). Additionally, we extracted intracranial volume measures using FreeSurfer v5.1 [76]. Briefly, this approach leverages the registration matrix between an individual image to a standard atlas space to estimate intracranial volume [77, 78].

#### c) Resting-state fMRI processing

The rs-fMRI data were processed as described previously [45] based on the configurable pipeline for the analysis of connectomes (C-PAC) (https://fcp-indi.github.io/) [79], including slice-time and head motion correction, skull stripping and intensity normalization. Data were corrected for head motion, as well as white matter and cerebrospinal fluid signals using CompCor [80], followed by band-pass filtering (0.01-0.1Hz). Both T1w and rs-fMRI data were linearly co-registered and mapped to MNI152 space. Functional imaging data were mapped to corresponding mid-thickness surfaces. We resampled data to the Conte69 surface template [75] via Workbench [74] with 10,242 surface points (vertices) per hemisphere. Due to the ongoing debate about global signal regression (GSR) [81], we did not apply GSR, but conducted additional control analyses using data that underwent GSR. We smoothed timeseries using a 5 mm full-width-at-half-maximum Gaussian kernel and computed intra-hemispheric functional connectivity (FC) as the Pearson correlation between all pairs of vertices within each hemisphere. FC matrices were Fisher r-to-z-transformed, to render correlation coefficients more normally distributed. Subjects with a mean framewise displacement (FD) >0.3 mm in rs-fMRI (two SD from the mean across all subjects) were excluded (n=9). Data were harmonized for site effects while preserving effects of age and ASD diagnosis using ComBat [82], which minimizes site-specific scaling factors by estimating their additive and multiplicative influence in a linear model, using empirical Bayes to predict site parameters more accurately [83, 84].

#### d) Connectivity distance profiling

We integrated FC and geodesic distance measures to compute CD profiles, as described previously [25, 43, 44]. Each participant’s functional connectivity matrix was masked to only consider the 10% strongest (*i*.*e*., highest absolute) values in each hemisphere. This threshold was chosen as prior literature suggested potential bias towards inter-individual differences with stricter cutoffs, and corresponding loss of functional specificity with more lenient thresholds [31]. Previous literature applying similar methodology applied a similar cutoff [43, 44]. CD was computed by binarizing thresholded functional connectivity matrices and retrieving the row-wise average geodesic distance in these nodes, generating a single value per vertex for each participant. The resulting CD maps, thus, reflects the average distance on the cortex from each region to the areas to which it is strongly connected to, thereby combining anatomical and functional information.

## Statistical analysis

We fit surface-based linear models correcting for age and head motion (as measured by mean framewise displacement) at each vertex *i*.

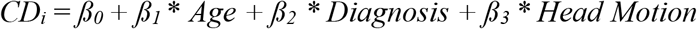

and group differences between ASD and NT were assessed in vertex-wise two-tailed Student’s t-tests using the BrainStat toolbox (https://github.com/MICA-MNI/BrainStat) [56]. Additionally, we computed vertex-wise effect sizes using *Cohen’s d*. Resulting p-values were adjusted for multiple comparisons using random field theory correction [85].

Several *post-hoc* analyses investigated the relationship between CD and behavioral metrics. Within ASD, we tested for associations to ASD symptom severity scores, specifically, total ADOS scores and ADOS subscores for communication, social interaction, and repetitive behaviors. In both ASD and NT groups, we examined the correlation to total IQ as well as verbal and performance IQ subscores, and the ratio of verbal and performance IQ [70]. First, we assessed the whole-brain association of CD with behavioral metrics in a vertex-wise linear model as described above. Additionally, we assessed the association to behavioral metrics within clusters of significant group differences which were identified in the previous step. Mean CD values were extracted from each cluster and the correlation to IQ and ADOS metrics were determined, adjusting for multiple comparisons using a false discovery rate (FDR) correction method [86].

### Contextualization to macro- and microscale principles of cortical organization

We explored effect sizes for group differences between ASD and NT individuals, *i*.*e*. Cohen’s *d*-values, and studied associations to macro- and microscale cortical patterns. For the whole-brain large-scale investigation, we analyzed mean CD and effect sizes in each of the seven previously described intrinsic functional networks [21]. Subsequently, we assessed the spatial correlation between Cohen’s *d* effect sizes and scores of the principal functional gradient that describes sensory-transmodal functional differentiation [24], while accounting for spatial autocorrelation with 5000 spin permutations [87]. We similarly investigated effect sizes within cytoarchitectonic cortical types of Von Economo and Koskinas [53, 54] leveraging the ENIGMA toolbox [57]. Finally, we assessed associations to the BigBrain histology gradient that describes microstructural differentiation [88], again sourced from the ENIGMA toolbox [57].

## RESULTS

### Reduced connectivity distance (CD) in autism

Mean CD in NT was higher within transmodal regions (*i*.*e*., heteromodal and paralimbic regions), such as the prefrontal and cingulate cortex and temporo-occipital-parietal junction, while CD values were lower in primary sensory and motor regions (**Figure 1A**).

**Figure 1.**
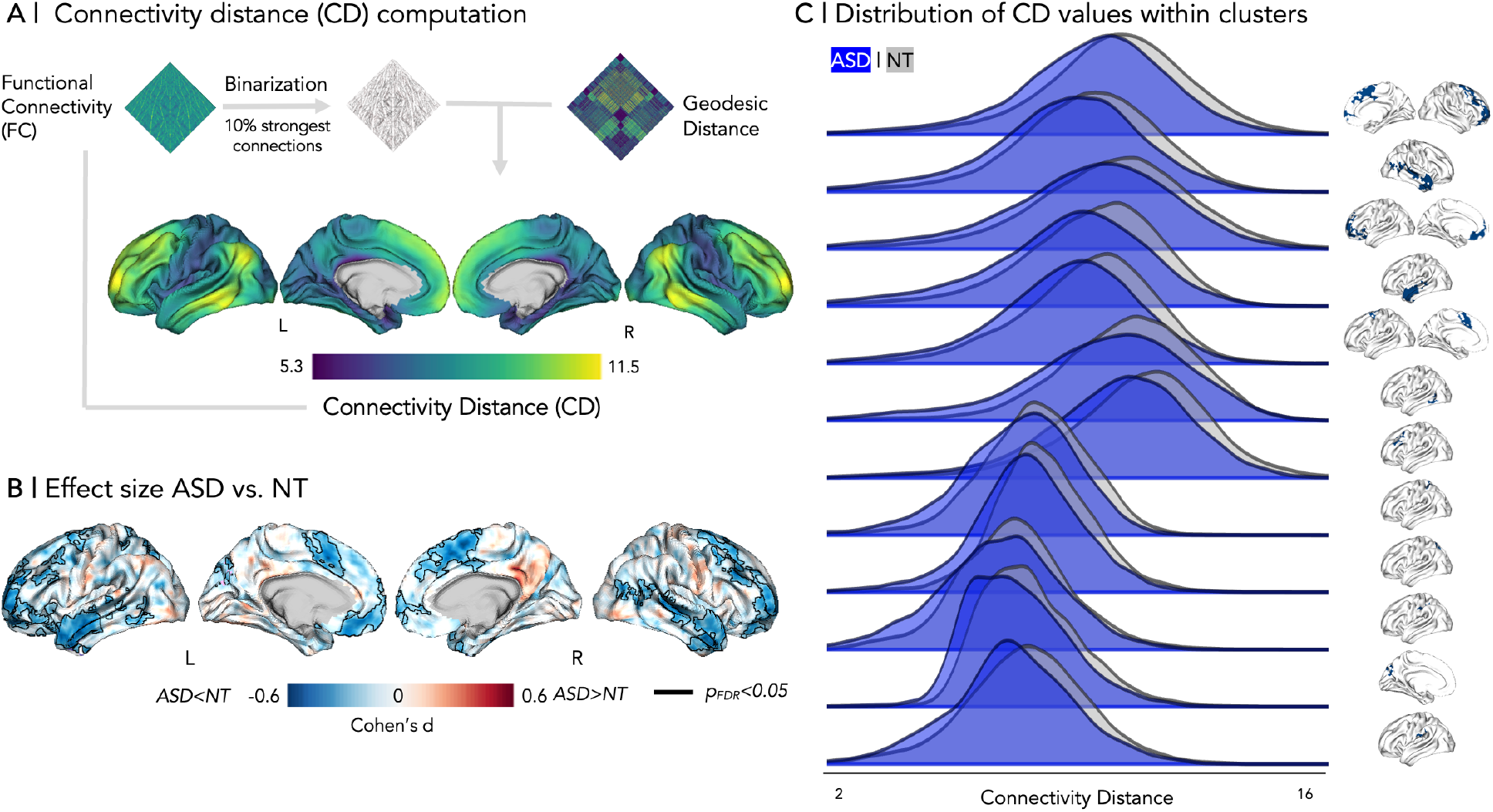
**A** | Workflow for CD computation and average CD maps. **B** | Effect size map for group differences (ASD *vs* NT). Clusters of significant changes after multiple comparisons correction are outlined (p_FDR_<0.05, vertex-based linear model). **C** | Distribution of CD values within clusters of significant reduction.

Comparing groups with linear models that additionally corrected for age and head motion revealed diffuse CD reductions in ASD relative to NT (**Figure 1B**), with twelve clusters deemed significant after multiple comparisons correction (*p*_*RFT*_<0.05). Clusters were mainly localized in the left and right temporal lobes and left prefrontal cortex (**Figure 1B, 2A**). *Cohen’s d* effect sizes for between-group differences amounted to *d=*0.600 in the temporal lobes as well as in the left prefrontal cortex (**Figure 1B**). Overall, there was a consistent shift of the CD distributions in ASD relative to NT across all significant clusters **(Figure 1C)**. Of note, CD was averaged per vertex, hence, the apparent decrease mirrors an increase in short-range connections at the expense of long-range links [43].

### Effects of age, symptom severity and intelligence measures

Within clusters of significant ASD-related reductions identified in the vertex-wise model, there was no significant correlation to age (*r*=0.151, *p*_*FDR*_=0.274, **Table 2)**. Moreover, there were no between-group effects when comparing children and adults, which indicated that ASD-related CD reductions were stable across pediatric and adult cohorts (*t*=-0.469, *p*_*FDR*_=0.640, **Figure 2B, Table 2**).

**Table 2.**
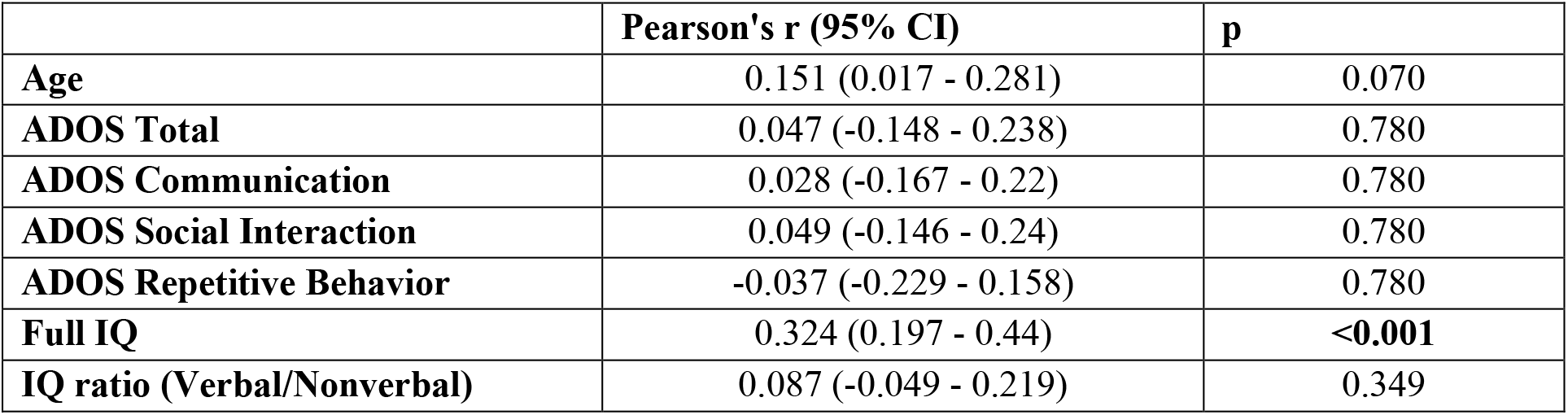
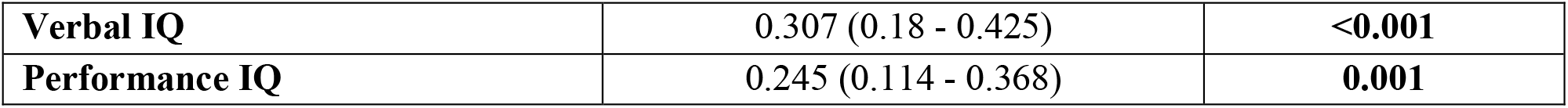
Correlation between mean CD (CD) in clusters of significant group differences and age, age group (Children <18 years and adults >= 18 years) as well as behavioral metrics. P-values were corrected for multiple comparisons using false discovery rate correction (q<0.05). Supplemental Table 1 provides values for each of the twelve clusters separately.

**Figure 2:**
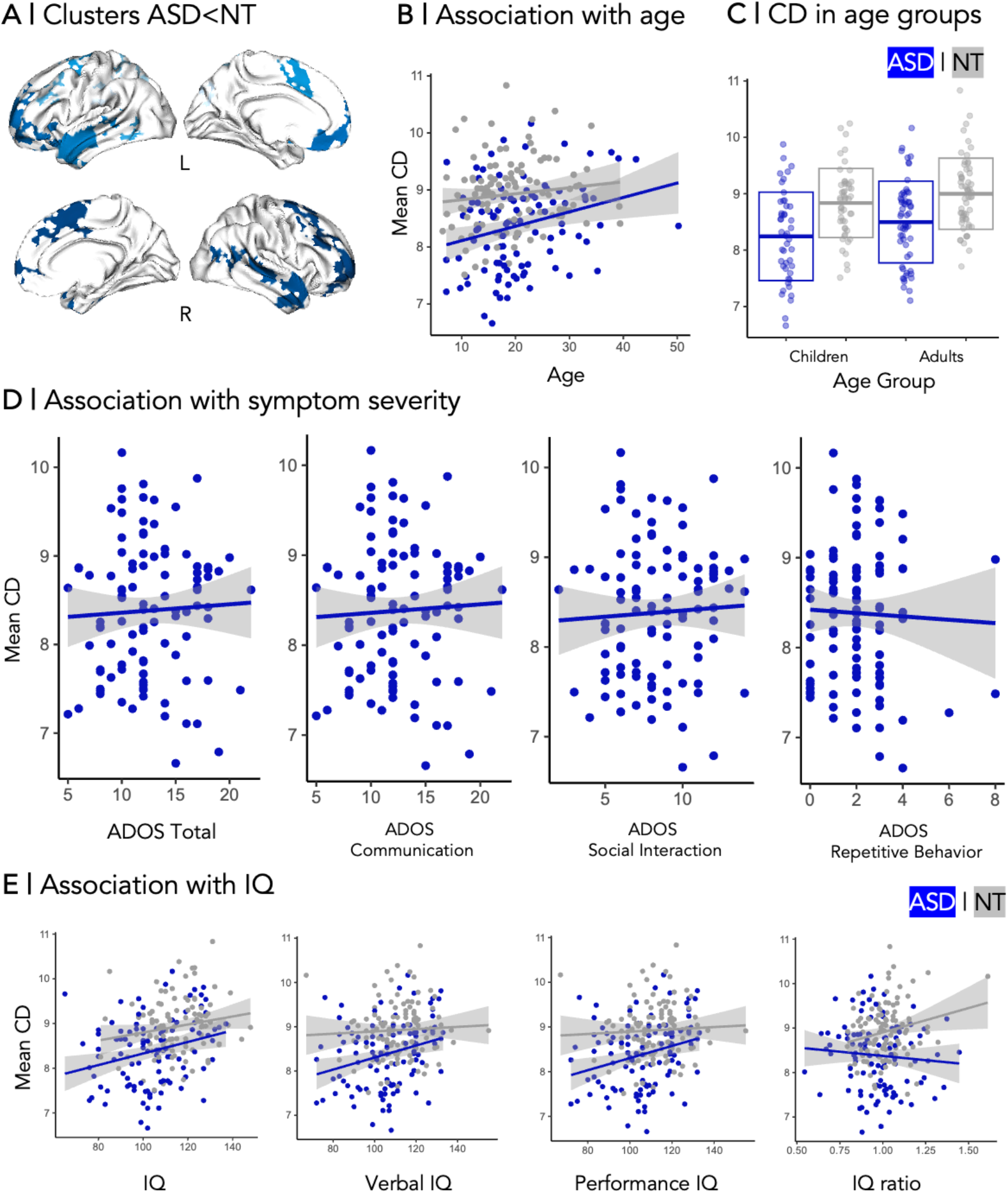
**A** | Clusters of significant CD reduction in ASD *vs* NT. **B-E** | Correlation between mean CD in clusters and age, age group (children <18 years, adults >18 years), total ADOS score, ADOS subscores for communication, social interaction, and repetitive behavior, as well as full IQ and IQ subscores for verbal and nonverbal IQ. IQ ratio denotes the ratio of verbal over nonverbal IQ. Correlation coefficients are listed in **Table 2**.

Within clusters of significant between-group differences, CD in the ASD group also did not show any significant associations with ADOS symptom severity scores (*r*=0.047, *p*_*FDR*_>0.4) nor with subscores for communication (*r*=0.028, *p*_*FDR*_>0.4), social interaction (*r*=0.049, *p*_*FDR*_=>0.4), and repetitive behaviors and interests (*r*=-0.037, *p*_*FDR*_>0.4). On the other hand, when averaged within all clusters, there was a significant association of CD with full (*r*=0.324, *p*_*FDR*_<0.001), verbal (*r*=0.304, *p*_*FDR*_<0.001), and performance IQ (*r*=0.245, *p*_*FDR*_=0.001; **Table 2**), while no significant interaction between diagnosis group and IQ measures was detectable (full IQ: *r*=-0.004, *p*_*FDR*_=0.561, verbal IQ: *r*=0.005, *p*_*FDR*_=0.561, performance IQ: *r*=-0.011, *p*_*FDR*_=0.214, IQ ratio: *r*=1.476, *p*_*FDR*_=0.128). Similar results were obtained in cluster-wise analysis. **Figure 2C-D** shows correlation between behavioral metrics and mean CD across all clusters, while separate cluster-wise analyses can be found in **Supplemental Figure 2 & Supplemental Table 1**.

In addition to cluster-wise analyses, we assessed whole-brain associations between CD and clinical metrics using a linear model of the influence of a behavioral score on CD while accounting for head motion and age. Overall, vertex-wise findings were consistent with within-cluster results (**Supplemental Figure 3**).

**Figure 3.**
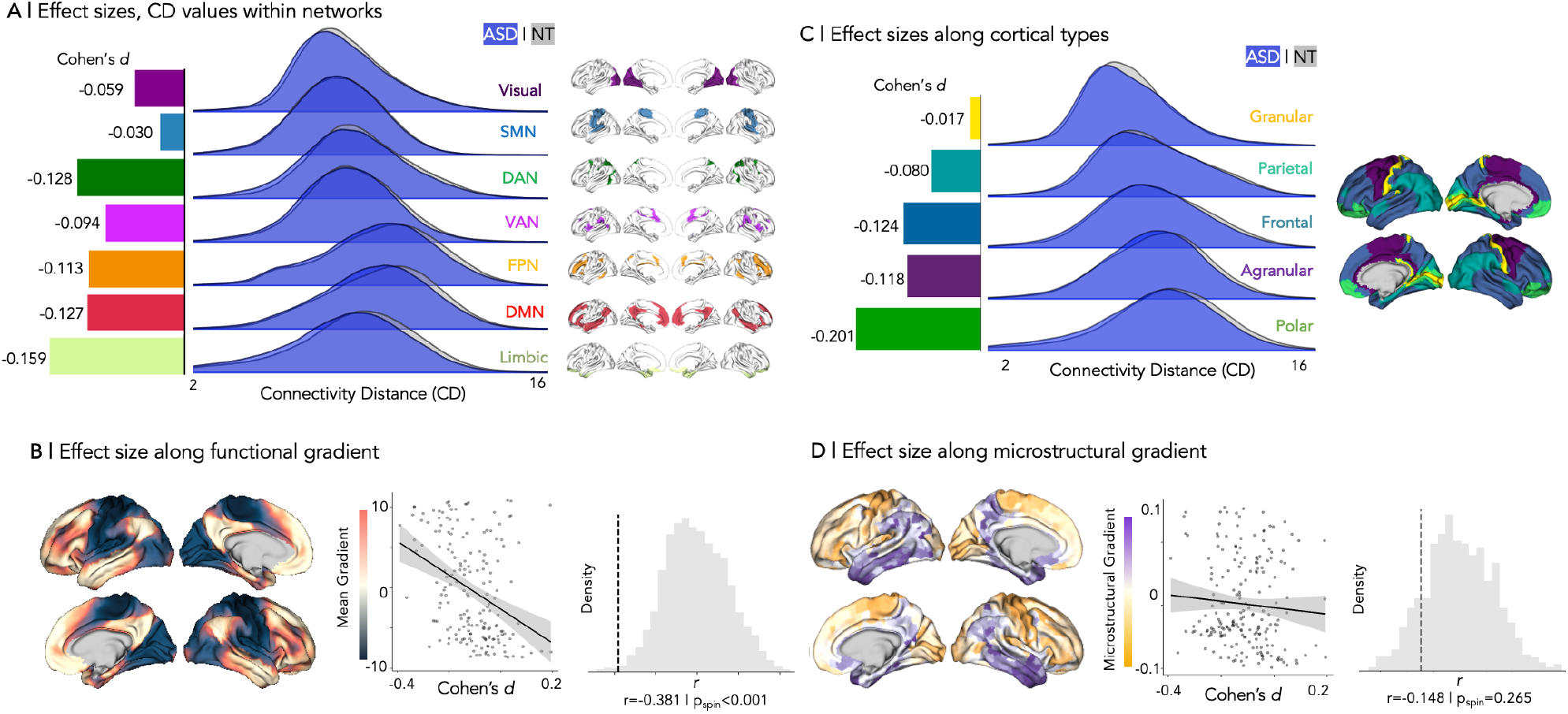
Relation to cortical organization. **A** | Effect sizes of between-group differences (ASD *vs* NT) in functional CD measures, stratified within seven intrinsic networks proposed by Yeo, Krienen, *et al*. [21] **B** | Correlation to the principal functional gradient [24]. Spatial associations were assessed using Spearman’s rank test, and p-values were adjusted for spatial autocorrelation using a spin test [87, 89]. **C** | Effect size for the between-group difference in functional CD in each cortical type as proposed by Von Economo and Koskinas [54]. **D** | Association to microstructural gradients derived from the 3D BigBrain [55]. *Abbreviations:* SMN = somatomotor network, DAN = dorsal attention network, VAN = ventral attention network, FPN = frontoparietal network, DMN= default mode network.

### Relation to cortical organization

#### Macroscale functional contextualization

Effect sizes differed across seven intrinsic functional networks [21], with higher effects towards transmodal compared to sensory/motor networks and peak reductions in CD in the limbic network (*Cohen’s d*=-0.159, **Figure 3A**). These findings were recapitulated when correlating effects against the intrinsic functional gradient [24] (*r=*-0.381, *p*_*spin*_*<*0.001), which were significant even when using models correcting for spatial autocorrelation (**Figure 3B**).

#### Microstructural and cytoarchitectonic contextualization

For cortical types as proposed by Von Economo and Koskinas [54], we were able to see notable effect size variations that also became larger towards limbic/paralimbic regions, with highest effect sizes being present in agranular polar cortex (*Cohen’s d*=0.201, **Figure 3C**). On the other hand, there was only a weak correlation between effects sizes and the primary BigBrain microstructural gradient (*r=*-0.148, *p*_*spin*_=0.265; **Figure 3D**).

## Control analyses

To ensure the robustness of our results across different data processing methods, we repeated analyses using functional connectivity that underwent GSR. **Supplemental Figure 2** shows respective effect size maps for both CD differences from uncorrected and from GSR-corrected functional CD maps. In both contrasts, highest between-group differences were localized in the left temporal lobe and left prefrontal cortex, as well as the right frontal lobe. However, group differences appeared higher for uncorrected data in medial prefrontal regions bilaterally, as well as the right temporal lobe. The congruence of both maps (*r*=0.546, *p*<0.001) suggested robustness of our results with respect to GSR application *vs* omission during data preprocessing. Additionally, we assessed intracranial volume as a potential confound for geodesic distance measures. There was no significant between-group difference in intracranial volume (*t=*0.694, *p*=0.488). Of note, there were eminent site-related differences in ADOS scores, suggesting behavioral phenotype differences between the cohorts or variations in clinical symptom severity assessment (*F=*3.904, *p=*0.023, **Supplemental Figure 4**).

## DISCUSSION

Atypical brain connectivity is thought to be at the core of autism [90] and our study examined whether ASD implicates long-range underconnectivity and local overconnectivity as compared to NT. We combined functional connectivity analysis with cortex-wide geodesic distance mapping and characterized overall shifts in the spatial extent of regional connectivity profiles [25, 43, 44]. Leveraging the open-access ABIDE dataset composed of ASD and NT individuals, we found robust evidence for global CD reductions in ASD, hinting towards a deficiency in long-range connections and a compensatory increase in short-range connectivity. Examining associations to functional topography, we noted a significant correlation to sensory-transmodal functional gradients, with most marked findings in paralimbic and heteromodal functional zones [24]. Likewise, contextualization against cytoarchitectural taxonomy [53, 54, 57] revealed most marked effects in agranular and polar cortices with low laminar differentiation. Findings were relatively robust irrespective of using GSR during preprocessing and were consistent across the different sites and age strata included in the study. Furthermore, CD reductions were stable across symptom severity metrics. While our findings did, thus, not capture imaging correlates of inter-individual differences in autism symptom load, they constitute a stable imaging phenotype of the condition. Moreover, CD alterations were positively correlated to both verbal and performance IQ measures, suggesting that these imbalances may index overall cognitive function in ASD. Collectively, imbalances in connectivity length distribution constitute a stable imaging phenotype of atypical neurodevelopment in ASD, representing a gradient of susceptibility closely intertwined with overarching principles of cortical organization.

The CD measure employed in this work combined rs-fMRI as an established *in vivo* proxy for functional interactions [91] with cortex-wide geodesic distance mapping [24, 43, 92, 93] to profile average functional connection length. This metric has been suggested to mirror functional and hierarchical properties of cortical areas, and to stratify systems in a data-driven, yet anatomically meaningful way [31, 32]. Prior research and our current findings have consistently demonstrated that long-range cortico-cortical connections predominate in transmodal networks, which comprise heteromodal association systems such as the DMN as well as paralimbic cortices [26, 39, 45]. These connections ensure efficient integration of functional signals in higher-order networks that are increasingly involved in abstract, integrative, as well as internally-generated cognitive and affective processes [16, 24, 94]. The observed decrease in long-range connectivity in ASD in this study may indicate brain reorganization characterized by reduced inter-network connectivity together with compensatory strengthening of local connections [41]. The topography of CD changes in ASD may indeed be particularly meaningful in the context of cortical hierarchies, a conjecture supported by the macro- and microscale contextualization analyses conducted in the current work. Here, we observed a sensory-fugal pattern of ASD-related CD alterations when cross-referencing our findings to both intrinsic functional and microstructural measures. Specifically, we observed that ASD-related CD alterations mirrored the intrinsic functional gradient of information abstraction, which runs from primary input to unimodal regions to higher-level cognition in transmodal cortices [30, 45]. Stratifying findings across seven intrinsic networks derived in previous work [21], we also found the largest effect sizes in limbic networks, confirming that ASD-related CD alterations primarily affect paralimbic/fugal systems. The latter conclusion could be supported by cross-referencing ASD-related effects to microstructure-based neural data. In fact, cytoarchitecture type stratification of our *in vivo* imaging-derived metrics revealed the strongest effect sizes in limbic agranular and polar cortices [53].

Our findings demonstrated increased vulnerability for ASD-associated connection length contractions in heteromodal and paralimbic cortices that collectively make up the transmodal core of cortical organization [62, 95-97]. When considering anatomical but also functional connectivity relationships across the cortex, both paralimbic as well as heteromodal association cortices are situated at a high distance from primary sensory and motor regions interacting with the here and now [98, 99]. When considering cortical microarchitecture, particularly its laminar organization, there has been evidence for an axis that differentiates sensory/motor on the one end from paralimbic systems on the other end [58, 60, 100, 101]. These characteristics are highlighted in spatial variations in the visibility of the internal granular layer, commonly referred to as layer IV [98, 102, 103]. In effect, the agranular cortex lacks the respective layer, the dysgranular cortex exhibits rudimentary layer IV characteristics, and the granular cortex shows clear layer IV [98]. Limbic and paralimbic cortical systems comprise mainly the agranular and dysgranular extent of this spectrum [98, 99, 104]. As such, paralimbic systems are microstructurally most segregated from granular systems interacting with the external environment [60]. Converging, but also somewhat different from the microstructural gradient is the sensory-association functional gradient, which radiates from sensory and motor systems towards heteromodal areas in the DMN [24, 60, 100]. These regions may contain complex microstructural signatures, including agranular types as well as granular cortices [23, 53, 54, 95]. In both heteromodal association systems as well as paralimbic cortices, there is prior evidence to suggest increased susceptibility to neurodevelopmental perturbations [58, 98]. Deficiency in long-range connectivity potentially results from cellular and laminar alterations [43], which may present a common substrate for different psychiatric and neurodevelopmental conditions [47, 58]. Risk genes for ASD as well as other neurodevelopmental conditions impact corticogenesis as early as in germinal stages [105, 106], impacting later axonal development that may disproportionally affect heteromodal systems [50, 107]. In future work, it remains thus to be established whether the current findings are specific to autism, or also visible in related neurodevelopmental indications. Our results, nevertheless, provide further justification to study microcircuit and macroscale alterations based on compact intermediary phenotypes such as CD. As such, they establish a perspective for future research, in order to better investigate and ultimately understand differentially impacted cortical hierarchies across common neurodevelopmental conditions.

Neuroimaging correlates of the typically developing connectome indicate marked shifts from local towards more distributed network patterns connections, while facilitating signaling across lobes and hemispheres [108, 109]. This progression highlights increased functional integration across brain networks, with short-range connections undergoing functional refinement [110, 111]. As such, network characteristics change from emphasized local processing in children to spatially and functionally distributed effects [109, 111, 112]. A potential microscale developmental mechanism driving this redistribution in typical development may be synaptic pruning [113-116]. In addition to its role in healthy brain maturation [117, 118], atypical pruning has been suggested in ASD [113, 119-121]. In effect, these alterations may be associated with local overconnectivity while not ensuring reliable long-range information relay [113, 119]. While speculative, our results potentially suggest a differential impact of pruning between long- and short-range connections, specifically, a higher requirement for longer connections to be retained in autism.

CD reductions identified in our work were invariant to age or symptom severity, possibly constituting a relatively stable marker of ASD-associated connectome reorganization. On the other hand, CD changes were modulated by intelligence measures, specifically, they exhibited an inverse correlation to IQ. Thus, our findings point towards contracted connectome profiles as markers for overall impaired cognitive performance. Conceptually, connectome contractions as imaged by CD implicate decreased communication efficiency [32, 43], offering a potential explanation for their potential contribution to general cognitive function in atypical development. Further research is needed to confirm this association in larger samples, and to also examine the specificity of these brain-behavior associations for autism *vis-à-vis* other neurodevelopmental indications.

