## Supplements for "Contracted Functional Connectivity Profiles in Autism"

### Supplementary Figures

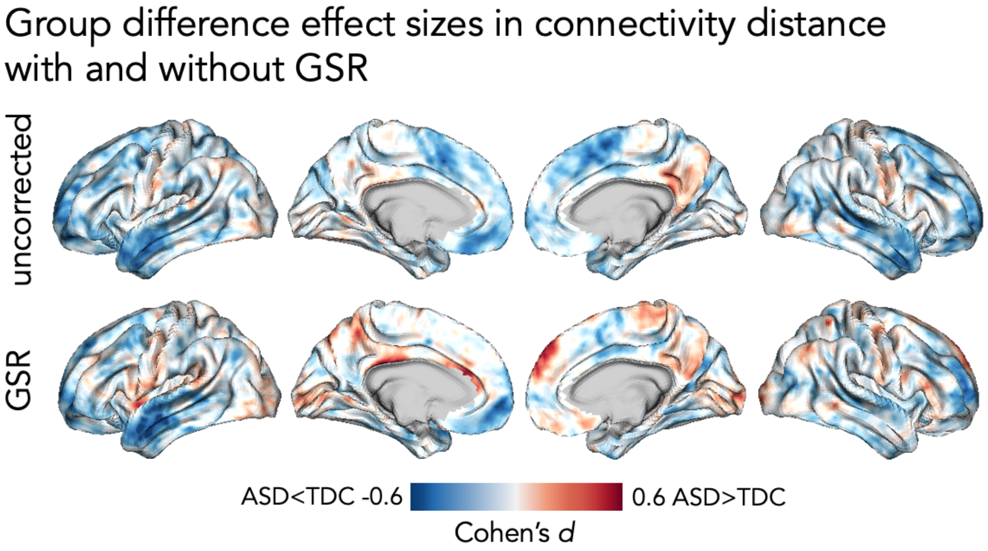


**Supplemental Figure 1 |** Effect size maps comparing CD between individuals with ASD and neurotypical controls. The bottom panel shows Cohen's d maps for connectivity maps based on functional connectivity that was corrected using global signal regression (GSR), while the top panel shows effect sizes in CD based on uncorrected functional connectivity.

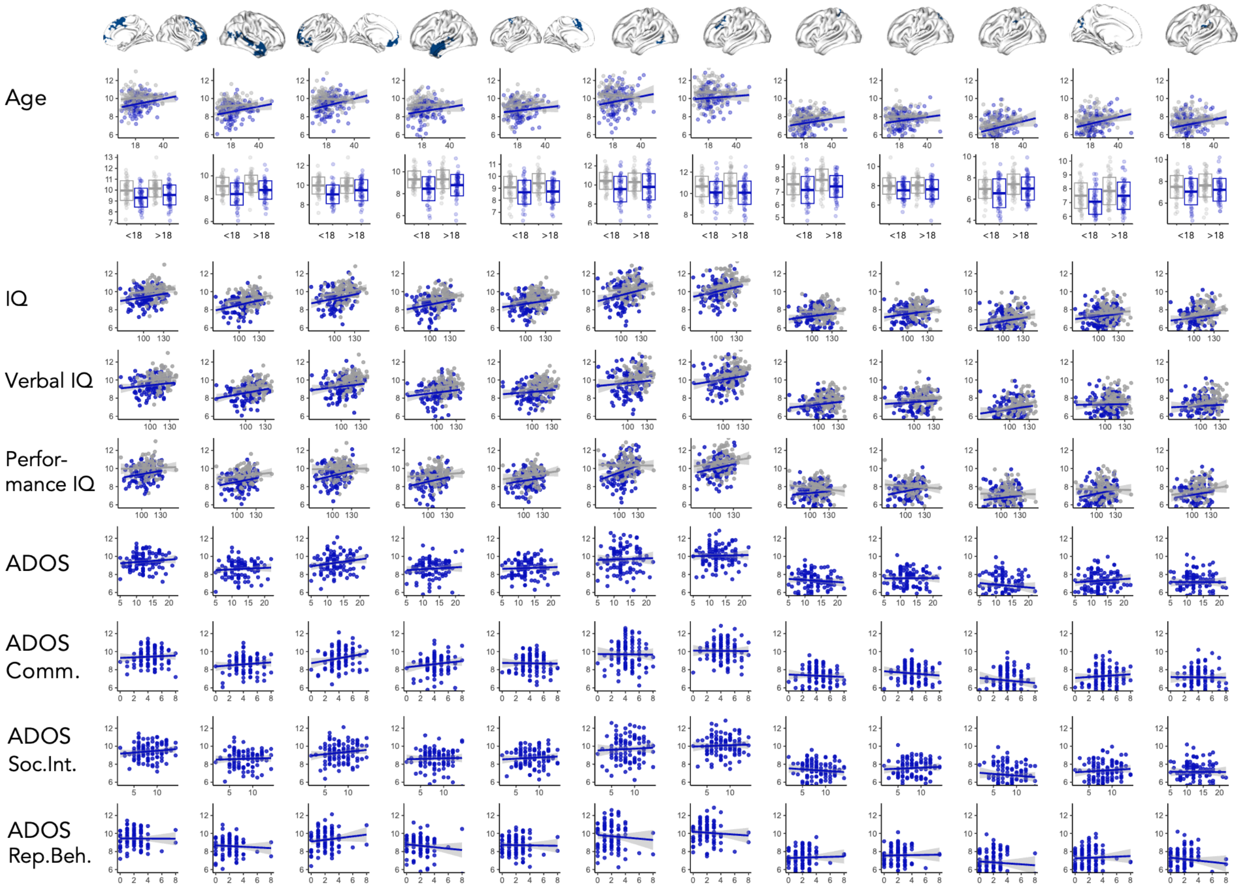


**SUPPLEMENTAL FIGURE 2 |** Correlation between CD and behavioral metrics for each cluster identified in a surface-based linear model separately. Respective correlation metrics can be found in supplemental table 1.

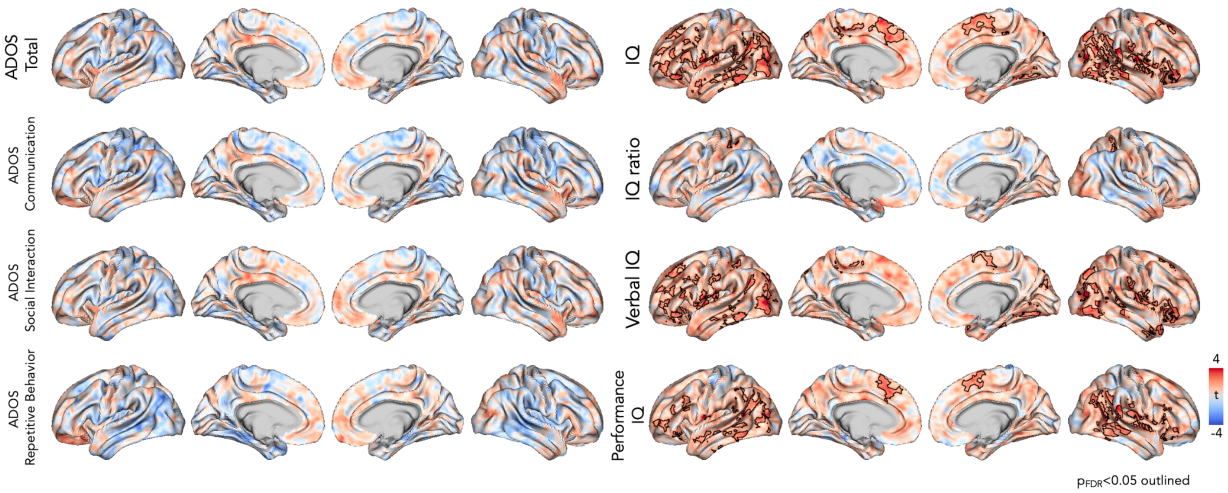


**Supplemental Figure 3 |** t-value maps for vertex-wise linear models assessing the influence of ADOS and IQ subscores on CD. Outlined clusters indicate areas of statistically significant CD increase with higher IQ metrics.

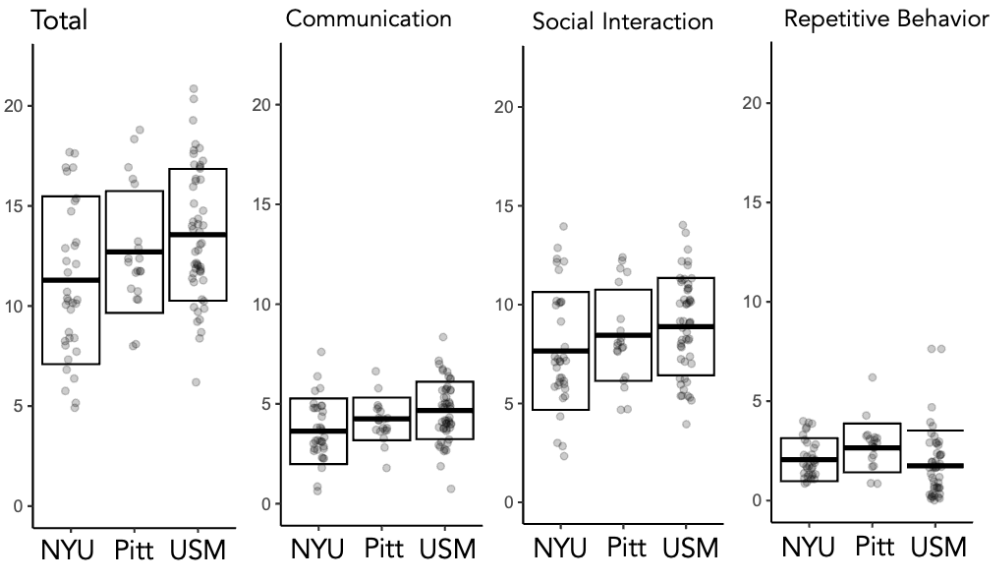


**SUPPLEMENTAL FIGURE 4 |** Variation of ADOS scores by site. NYU = New York University Langone Medical Center, Pitt = Pittsburgh School of Medicine, USM = University of Utah School of Medicine.

#### SUPPLEMENTARY TABLES

| | Age | | Age Group | |
|---|---|---|---|---|
| **Cluster** | **Pearson's r (95% CI)** | **p** | **Coefficient (95% CI)** | **p** |
| 1 | 0.133 (-0.002 - 0.263) | 0.142 | 0.061 (-0.013 - 0.134) | 0.251 |
| 2 | 0.143 (0.008 - 0.273) | 0.105 | 0.084 (0.008 - 0.161) | 0.089 |
| 3 | 0.093 (-0.043 - 0.225) | 0.372 | 0.048 (-0.021 - 0.117) | 0.367 |
| 4 | 0.068 (-0.067 - 0.202) | 0.557 | 0.04 (-0.03 - 0.109) | 0.483 |
| 5 | 0.116 (-0.019 - 0.248) | 0.222 | 0.055 (-0.016 - 0.127) | 0.292 |
| 6 | 0.022 (-0.113 - 0.157) | 0.873 | 0.006 (-0.051 - 0.063) | 0.921 |
| 7 | 0.015 (-0.12 - 0.15) | 0.921 | 0.002 (-0.06 - 0.063) | 0.978 |
| 8 | 0.168 (0.034 - 0.297) | 0.051 | 0.085 (0.014 - 0.157) | 0.062 |
| 9 | 0.083 (-0.053 - 0.215) | 0.454 | 0.023 (-0.051 - 0.098) | 0.718 |
| 10 | 0.207 (0.074 - 0.333) | **0.013** | 0.088 (0.028 - 0.149) | **0.019** |
| 11 | 0.195 (0.062 - 0.322) | **0.020** | 0.096 (0.028 - 0.163) | **0.024** |
| 12 | 0.12 (-0.015 - 0.251) | 0.205 | 0.029 (-0.032 - 0.091) | 0.557 |
| | **ADOS Total** | | **ADOS Communication** | |
| **Cluster** | **Pearson's r (95% CI)** | **p** | **Pearson's r (95% CI)** | **p** |
| 1 | 0.111 (-0.084 - 0.298) | 0.483 | 0.056 (-0.14 - 0.246) | 0.758 |
| 2 | 0.069 (-0.127 - 0.259) | 0.684 | 0.095 (-0.101 - 0.283) | 0.557 |
| 3 | 0.183 (-0.011 - 0.364) | 0.165 | 0.202 (0.009 - 0.381) | 0.11 |
| 4 | 0.074 (-0.121 - 0.264) | 0.653 | 0.127 (-0.069 - 0.312) | 0.409 |
| 5 | 0.047 (-0.148 - 0.239) | 0.784 | -0.016 (-0.209 - 0.178) | 0.944 |
| 6 | 0.032 (-0.163 - 0.224) | 0.873 | -0.009 (-0.202 - 0.185) | 0.975 |
| 7 | 0.027 (-0.167 - 0.22) | 0.895 | -0.01 (-0.203 - 0.184) | 0.975 |
| 8 | -0.089 (-0.277 - 0.107) | 0.573 | -0.052 (-0.243 - 0.143) | 0.772 |
| 9 | 0.004 (-0.189 - 0.198) | 0.978 | -0.094 (-0.282 - 0.101) | 0.557 |
| 10 | -0.099 (-0.287 - 0.096) | 0.557 | -0.088 (-0.277 - 0.107) | 0.573 |
| 11 | 0.095 (-0.101 - 0.283) | 0.557 | 0.08 (-0.116 - 0.269) | 0.625 |
| 12 | -0.007 (-0.2 - 0.187) | 0.978 | -0.01 (-0.203 - 0.184) | 0.975 |
| | **ADOS Social Interaction** | | **ADOS Repetitive Behaviors** | |
| **Cluster** | **Pearson's r (95% CI)** | **p** | **Pearson's r (95% CI)** | **p** |
| 1 | 0.123 (-0.072 - 0.309) | 0.427 | -0.006 (-0.2 - 0.187) | 0.978 |
| 2 | 0.041 (-0.153 - 0.233) | 0.821 | -0.063 (-0.253 - 0.132) | 0.718 |
| 3 | 0.14 (-0.055 - 0.324) | 0.35 | 0.139 (-0.056 - 0.324) | 0.35 |
| 4 | 0.031 (-0.164 - 0.223) | 0.875 | -0.112 (-0.299 - 0.083) | 0.483 |
| 5 | 0.075 (-0.12 - 0.265) | 0.651 | -0.019 (-0.211 - 0.176) | 0.929 |
| 6 | 0.049 (-0.145 - 0.241) | 0.783 | -0.075 (-0.265 - 0.12) | 0.651 |
| 7 | 0.044 (-0.151 - 0.235) | 0.809 | -0.073 (-0.263 - 0.122) | 0.653 |
| 8 | -0.094 (-0.282 - 0.102) | 0.557 | 0.032 (-0.162 - 0.224) | 0.873 |
| 9 | 0.06 (-0.135 - 0.25) | 0.729 | 0.02 (-0.175 - 0.212) | 0.928 |
| 10 | -0.088 (-0.276 - 0.108) | 0.573 | -0.064 (-0.254 - 0.131) | 0.717 |
| 11 | 0.086 (-0.109 - 0.275) | 0.577 | 0.048 (-0.146 - 0.24) | 0.783 |
| 12 | -0.003 (-0.197 - 0.19) | 0.978 | -0.108 (-0.296 - 0.087) | 0.499 |
| | **Full IQ** | | **IQ ratio** | |
| **Cluster** | **Pearson's r (95% CI)** | **p** | **Pearson's r (95% CI)** | **p** |
| 1 | 0.256 (0.125 - 0.378) | **0.002** | 0.071 (-0.065 - 0.204) | 0.543 |
| 2 | 0.329 (0.203 - 0.445) | **<0.001** | 0.098 (-0.037 - 0.23) | 0.349 |
| 3 | 0.301 (0.173 - 0.419) | **<0.001** | 0.105 (-0.03 - 0.237) | 0.292 |
| 4 | 0.285 (0.156 - 0.405) | **<0.001** | 0.061 (-0.074 - 0.195) | 0.573 |
| 5 | 0.264 (0.134 - 0.385) | **0.001** | 0.028 (-0.108 - 0.162) | 0.825 |
| 6 | 0.259 (0.129 - 0.381) | **0.002** | 0.016 (-0.119 - 0.151) | 0.921 |
| 7 | 0.292 (0.163 - 0.41) | **<0.001** | 0.079 (-0.057 - 0.212) | 0.483 |

| | | | | |
|---|---|---|---|---|
| 8 | 0.178 (0.044 - 0.306) | **0.037** | 0.169 (0.035 - 0.297) | 0.051 |
| 9 | 0.166 (0.032 - 0.295) | 0.053 | 0.038 (-0.098 - 0.172) | 0.758 |
| 10 | 0.189 (0.056 - 0.316) | **0.024** | 0.146 (0.011 - 0.276) | 0.097 |
| 11 | 0.154 (0.019 - 0.283) | 0.081 | -0.034 (-0.168 - 0.102) | 0.783 |
| 12 | 0.199 (0.066 - 0.325) | **0.018** | 0.002 (-0.133 - 0.137) | 0.978 |
| | **Verbal IQ** | | **Performance IQ** | |
| **Cluster** | **Pearson's r (95% CI)** | **p** | **Pearson's r (95% CI)** | **p** |
| 1 | 0.19 (0.056 - 0.317) | **0.024** | 0.245 (0.114 - 0.368) | **0.003** |
| 2 | 0.242 (0.11 - 0.365) | **0.003** | 0.316 (0.189 - 0.432) | **<0.001** |
| 3 | 0.212 (0.079 - 0.337) | **0.01** | 0.296 (0.167 - 0.414) | **<0.001** |
| 4 | 0.226 (0.094 - 0.35) | **0.007** | 0.26 (0.13 - 0.382) | **0.002** |
| 5 | 0.215 (0.082 - 0.34) | **0.009** | 0.229 (0.097 - 0.353) | **0.006** |
| 6 | 0.229 (0.097 - 0.353) | **0.006** | 0.225 (0.092 - 0.349) | **0.007** |
| 7 | 0.222 (0.089 - 0.346) | **0.007** | 0.28 (0.151 - 0.4) | **<0.001** |
| 8 | 0.067 (-0.069 - 0.2) | 0.557 | 0.221 (0.088 - 0.345) | **0.007** |
| 9 | 0.143 (0.008 - 0.273) | 0.105 | 0.152 (0.017 - 0.281) | 0.083 |
| 10 | 0.092 (-0.043 - 0.225) | 0.373 | 0.223 (0.09 - 0.347) | **0.007** |
| 11 | 0.153 (0.018 - 0.282) | 0.082 | 0.116 (-0.019 - 0.248) | 0.222 |
| 12 | 0.179 (0.045 - 0.307) | **0.035** | 0.166 (0.032 - 0.294) | 0.053 |

**SUPPLEMENTAL TABLE 1 |** Correlation between CD and age and behavioral metrics within each cluster that reached statistical significance in a surface-based linear model. P-values were adjusted for multiple comparisons using false discovery rate correction ($q<0.05$).
